# Intermittent treatment with elamipretide preserves exercise tolerance in aged female mice

**DOI:** 10.1101/2022.11.29.518431

**Authors:** Matthew D. Campbell, Ashton T. Samuelson, Ying Ann Chiao, Mariya T. Sweetwyne, Warren C. Ladiges, Peter S. Rabinovitch, David J. Marcinek

## Abstract

The pathology of aging impacts multiple organ systems including the kidney, skeletal, and cardiac muscle. Long-term treatment with the mitochondrial targeted peptide elamipretide has previously been shown to improve *in vivo* mitochondrial function in aged mice that is associated with increased fatigue resistance and treadmill performance, improved cardiovascular diastolic function, and glomerular architecture of the kidney. However, elamipretide is a short tetrameric peptide that is not orally bioavailable limiting its routes of administration. This study tested whether twice weekly intermittent injections of elamipretide could recapitulate the same functional improvements as continuous long-term infusion. We found that intermittent treatment with elamipretide for 8 months preserved endurance running in mice, skeletal muscle force production, and left ventricular mass but did not affect heart or kidney function as previously reported using continuous treatment.

## 1. Introduction

Mitochondria play a key role in the biology of aging and age-related mitochondrial dysfunction contributes to age-related cellular, tissue, and end-organ degeneration, thereby increasing the susceptibility to age-related chronic disease. This is particularly relevant in tissues with a high energetic demand such as skeletal and cardiac muscle, which contributes significantly to the loss of mobility (*1*), exercise intolerance (*2*) and an associated decline in quality of life (*3, 4*). As a result of the important role of mitochondrial-related energy production in aging heart and skeletal muscle, there is increasing interest in mitochondrial-targeted interventions that have the potential to reverse and slow the decline in sarcopenia, cardiac dysfunction, exercise intolerance and reduced healthspan. Previous work has demonstrated that the mitochondrial-targeted peptide elamipretide (ELAM), previously referred to as Bendavia, MTP-131, and SS-31, is an effective intervention to reverse skeletal muscle (*5, 6*), cardiac (*7*), kidney (*8*), retina (*9*), and cognitive dysfunction in aged mice (*10*). Clinical trials in aged humans demonstrated improvements of *in vivo* mitochondrial energetics after a single treatment (*11*). This improvement was short-lived, consistent with the short half-life of the compound(*5*).

In order to compare the ability of ELAM to slow aging or increase lifespan to other more established aging interventions an extended duration of treatment is necessary. Aging remains the greatest risk factor for the development of multiple diseases, including heart disease (*12, 13*), cancer (*14*), and sarcopenia (*15*). Therefore, slowing the aging process has the potential to improve public health, increase life expectancy, and expand the years over which people can expect to live independent and productive lives. There are two strategies for testing compounds that can improve function with age. The first is to test whether a compound is capable of reversing age-related decline when administered late in life, after the effects of age are already apparent. The second strategy is to determine whether or not an intervention can slow the aging process when the agent is started early, before significant dysfunction is present, and then continued with advancing age.

Currently, ELAM is not available as an oral formulation because of its poor oral bioavailability. However, long-term studies in humans evaluating the effects of ELAM for healthspan and longevity have successfully used daily, subcutaneous doing. While this route of administration has been shown to be effective, we sought to investigate whether or not an ELAM regimen 2x/week for 8 months would reproduce or improve the beneficial effects we have observed in the skeletal and cardiac function of aged mice with 8-weeks of continuous treatment.

## 2. Methods

### 2.1 Animals

All experiments performed in this study were reviewed and approved by the University of Washington Institutional Animal Care and Use Committee (IACUC). Female C57Bl/J mice were procured from the National Institute of Aging aged mouse colony. Mice were kept at 21°C on a 14/10 light dark cycle and given chow and water *ad lib*.

### 2.2 Elamipretide administration

Animals were weighed once per week to determine injection volumes for the week. Mice were given a twice weekly dose on Monday and Thursday of either saline or 3mg/kg of ELAM in saline via intraperitoneal injection for 8 months.

### 2.3 Treadmill performance

Mice were tested at baseline prior to initiation of twice weekly injections and every 4 weeks thereafter. All mice were acclimated to the treadmill twice prior to each endurance test. During acclimation day 1, animals were loaded onto the treadmill at 0° incline with an active shockplate delivering 0.15 amps at 1Hz, mice were allowed to free roam for 2 minutes and then run for 2 minutes, accelerating from 0 meters(m)/minute(min) to 10 m/min, then held at a constant speed of 10 m/min. During acclimation day 2, animals underwent the same steps as day 1 with a maximum speed of 20 m/min. Finally, on test days mice were placed on a treadmill at 10° incline and run for 5 minutes accelerating from 0 m/min to 30 m/min, then run to exhaustion at 30 m/min. Exhaustion was defined as inability to remount the treadmill while receiving 5 consecutive shocks plus light physical prodding. Treadmill acclimation and endurance tests were all performed between 8 pm and 2 am to align with the natural active period for these mice.

### 2.4 In vivo muscle mechanics and analyses

Mice were tested at baseline prior to initiation of 2X per week injections and at 8, 16, and 28 weeks during treatment. Animals were anesthetized using 2-4% isoflurane in 1 L/min O_2_ and maintained on a heated platform maintained at 37°C using a thermally controlled water circulator. The right hind leg was held in position at the kneecap and the foot taped to a servomotor arm for torque force measurement and linear conversion (Aurora Scientific, Aurora, ON, Canada). Stimulation was controlled by and measured using Dynamic Muscle Control software v.5.500 (Aurora Scientific). For fatigue tests the gastrocnemius (GAS) muscle was stimulated using a S88X dual channel stimulator (Grass Astro-Med Inc.) through the tibial nerve at 180 Hz every other second for 2 minutes followed by a 1 minute and 5 minute post-fatigue tetanic test to evaluate recovery. Force and fatigue analysis was performed using Dynamic Muscle Analysis software v. 5.300 (Aurora Scientific).

### 2.5 Skeletal and cardiac muscle respirometry and peroxide production

Immediately following cervical dislocation, GAS and heart were dissected and placed on ice. Approximately 2-3 mg of the red GAS was removed, teased apart, and permeabilized using 50 ug/mL saponin at 4°C for 40 minutes. During permeabilization approximately 10 mg of the apex of the heart was removed and homogenized. Permeabilized fibers and heart homogenate were added to a 2 mL chamber of an Oxygraph 2K dual respirometer/fluorometer (Oroboros Instruments, Innsbruck, Austria) at 37°C and stirred at 750 rpm during titrations of substrates, uncouplers, and inhibitors. Following addition of sample superoxide dismutase (5 u/mL), amplex red (10 uM), and horseradish peroxidase (0.5 u/mL) were added to the chambers. For evaluation of forward electron flow, titrations were performed in the following order (concentrations): malate (5 mM), pyruvate (5 mM), and glutamate (10 mM); sub-saturating ADP (50 uM), saturating ADP (2.5 mM), succinate (20 mM), rotenone (0.5 uM), antimycin A (2.5 uM), TMPD (0.5 mM) and ascorbate (2 mM), KCN (1 mM). For evaluation of reverse electron flow, titrations were performed in the following order (concentrations): succinate (1, 5, 10, and 20 mM), and saturating ADP (2.5 mM).

### 2.6 Echocardiography

Echocardiography was performed as previously described (*7*). In brief, animals were anesthetized using 0.5-1% isofluorane delivered via 1 L/min O2. Imaging was performed using a Siemens Acuson CV-70 equipped with a 13MHz probe. Results were analyzed using mixed-effects ANOVA with correction for multiple comparison’s using Sidak’s post hoc test.

### 2.7 Kidney Histology and Stoichiometry

At sacrifice the right kidney from each animal was decapsulated, bifurcated and fixed at 4°C in 4% paraformaldehyde for 16 hours followed by 70% ethanol before paraffin embedding. Immunohistochemistry for podocyte p57 expression (rabbit anti-p57; Santa Cruz Biotechnology, Santa Cruz, CA) was performed on 4 uM sections using antigen-retrieval EDTA buffer pH 6 and overnight incubation at 1:800. Antibody was visualized with diaminobenzidine (Sigma) and counterstained with Periodic Acid Schiff stain (PAS), including hematoxylin to visualize nuclei. At least 40 glomeruli from the outer cortex and 20 glomeruli from the juxtamedullary cortex were scored from each kidney (n=5 ELAM treated, n=4 control) for the presence of sclerosis on a scale of: 0 (no injury); 1 (minor structural changes or matrix deposition); 2-3 (25%-75% sclerotic); 4 (fully sclerosed or necrotic). Podocytes were measured in micrographs using ImageJ software and the Venkatereddy-Wiggins method (*16*) to calculate the number, diameter and density within the tuft for each glomerulus (n=3 ELAM treated, n=2 control). Individual student’s t-test showed no significant differences between ELAM and control for any kidney parameter.

## 3. Results

### 3.1 Body and tissue mass

Starting weights for the saline and ELAM-treated groups were not significantly different (28.3 g ± 1.7 g vs. 28.9 g ± 2.2 g). Two times per week treatment with ELAM starting at 20 months of age had no effect on body mass throughout the 8-month treatment period, analyzed by area under the curve (Fig. 1A), nor was there a difference in end of treatment weights. There was no effect of treatment on survival, with 8 saline mice and 9 ELAM mice dying of natural causes over the treatment period (Fig. 1B). Due to the small sample sizes, it was not possible to make any statements regarding differences in cause of death. Consistent with no differences in body mass, there was also no effect of treatment on skeletal muscle mass of the GAS and the tibialis anterior (TA) muscles and no effect on cardiac hypertrophy (Fig. 1 C-E).

**Figure 1.**
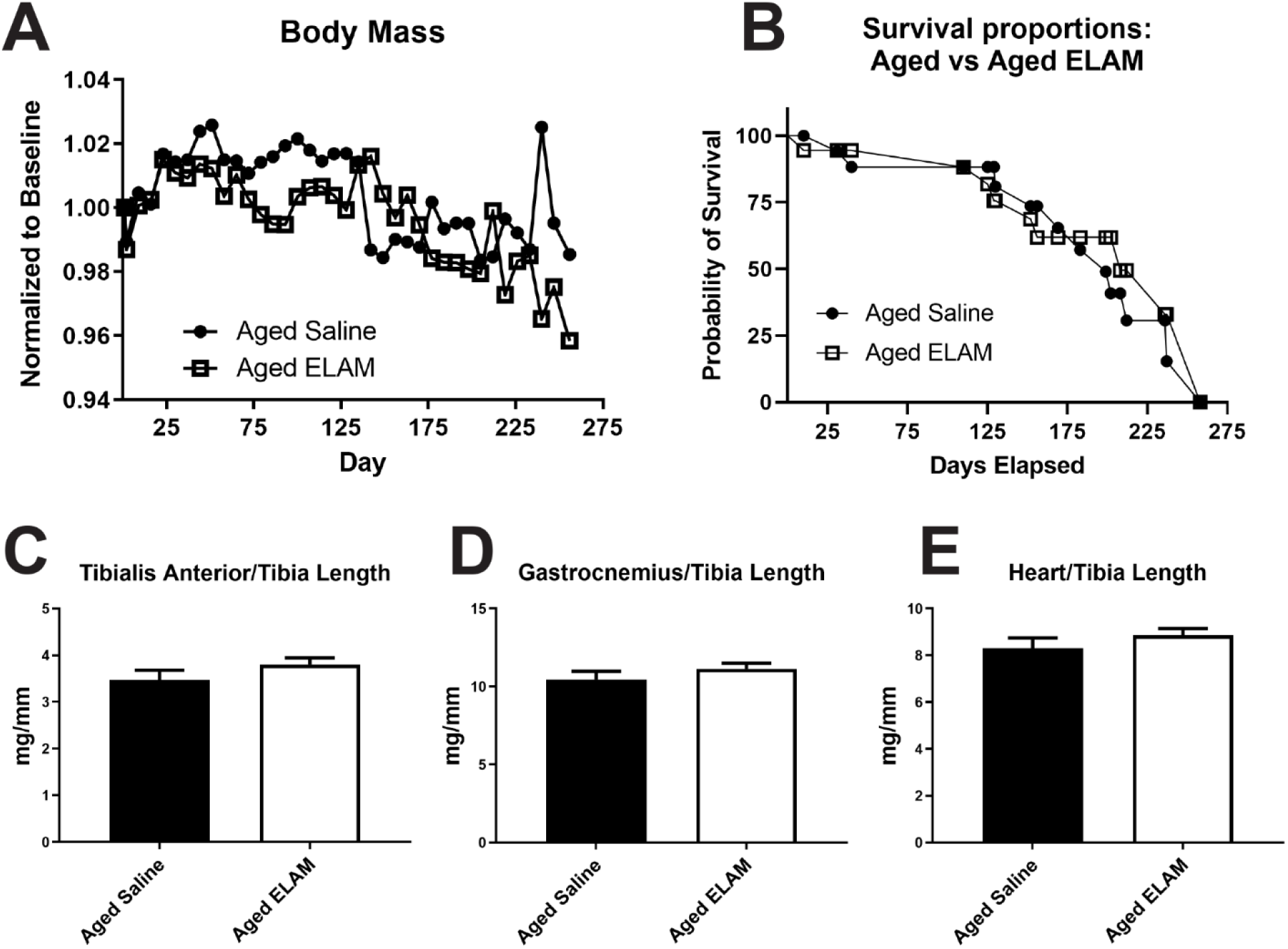
Mass and survival. **(A)** Weekly masse over the course of treatment normalized to initial body masses **(B)** Survival curves of animals that did not reach endpoint **(C)** Tibialis anterior mass normalized to tibia lenght **(D)** Gastrocnemius mass normalized to tibia length **(E)** Heart mass normalized to tibia length. Data expressed as means ± SE

### 3.2 Exercise tolerance

To assess the effect of treatment and age on whole body performance we tested exercise tolerance by running mice to exhaustion on a treadmill every 4 weeks. There was no difference in running time at baseline between the groups. With a mixed effect ANOVA analysis there was a highly significant effect of both age and treatment on total running time. The improvement in exercise tolerance was apparent after the first 4 weeks of treatment with ELAM and persisted throughout the 28-week treatment period (Fig. 2).

**Figure 2.**
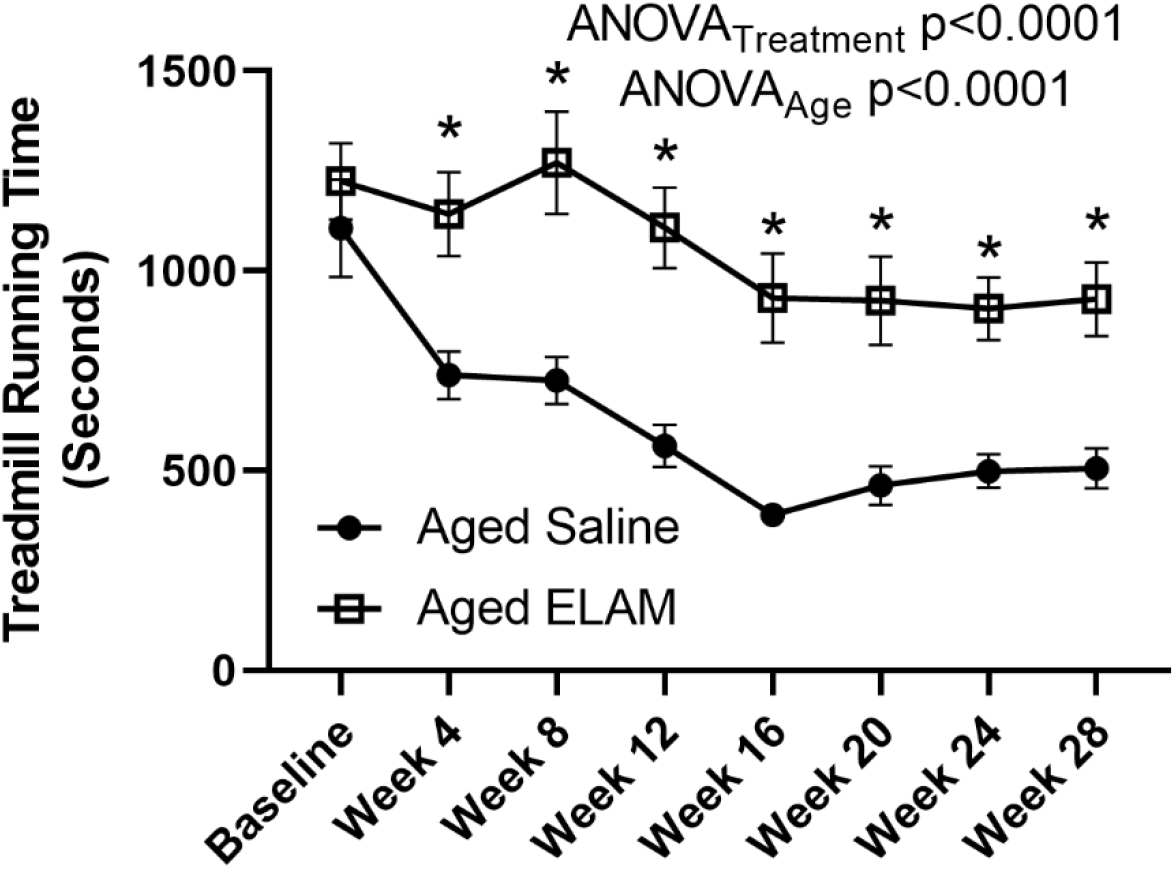
Treadmill running times. Elamipretide significantly preserves treadmill endurance. Data expressed as means ± SE. Mixed effects ANOVA, with Sidak’s correction for multiple comparisons, p<0.05.

### 3.3 Skeletal muscle and cardiac function

To assess the effect of aging and treatment on skeletal muscle function we measured the maximum force-time integral and fatigue resistance in the GAS muscle every 8 weeks throughout the treatment period. Despite the improved treadmill exercise tolerance in the ELAM group and previous work demonstrating increased fatigue resistance after continuous 8-week treatment (*6*), there was no effect of treatment on GAS fatigue resistance nor force recovery following fatigue at any time point. Fatigue curves for baseline and week 28 are shown in Fig. 3 A and B. There was also no significant effect of age on the maximum FTI of the GAS throughout the study period (p=0.07). Although the effect was not significant with a mixed model ANOVA, multiple testing revealed a borderline significant improvement in maximum FTI at week 16 only in the ELAM group (Fig. 3C).

**Figure 3.**
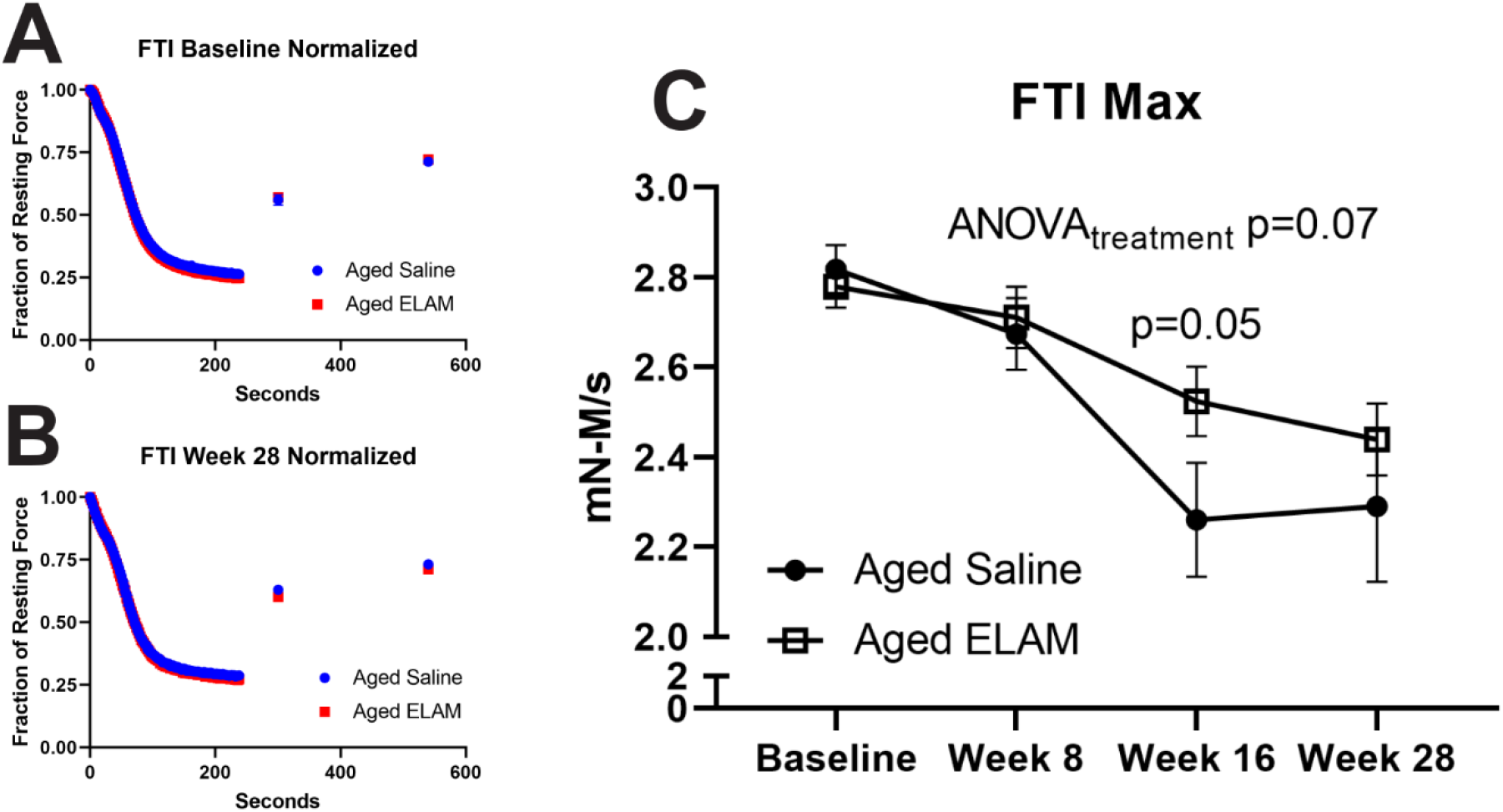
In vivo skeletal muscle function. **(A)** Force time integral of fatigue normalized to resting force at baseline before entering study **(B)** Force time integral of fatigue normalized to resting force at endpoint of study **(C)** Maximum force time integral over the course of the study. Data expressed as means ± SE. Mixed effects ANOVA, with Sidak’s correction for multiple comparisons, p<0.05.

To assess whether the 28-week intermittent treatment reproduces the reversal of cardiac dysfunction we performed echocardiography on the mice at baseline, 16, 24, and 28 weeks. We have previously reported that ELAM treatment reverses diastolic dysfunction in aged mice (*7*). Similar to the maximum FTI analysis, the long-term intermittent ELAM treatment pointed toward a non-significant treatment effect (Fig. 4A) p=0.07) with multiple testing indicating a significantly reduced Ea/Aa ratio at 28 weeks only (Fig. 4B). In contrast, the left ventricular mass index (LVMI) indicated a significant effect of treatment overall with no significant effect at any one timepoint p<0.01.

**Figure 4.**
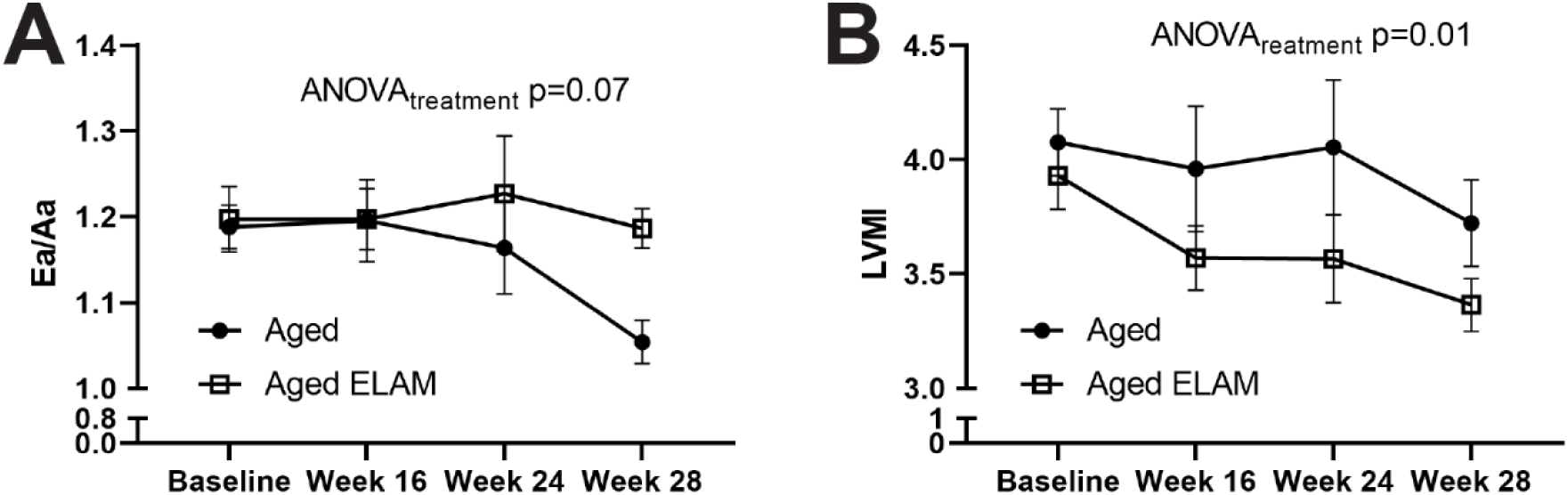
Cardiac effects. **(A)** Ea/Aa ratio over the course of the study **(B)** Left Ventricular Mass Index over the course of the study. Data expressed as means ± SE. Mixed effects ANOVA, with Sidak’s correction for multiple comparisons, p<0.05.

### 3.4 Mitochondrial function

To test the effect of long-term intermittent treatment of ELAM on mitochondrial function and mitochondrial peroxide production we analyzed permeabilized muscle fibers and heart homogenate using a combination fluorometer/respirometer (Oroboros Instruments, Innsbruck AU) at the end of 28 weeks. We tested forward electron flow and reverse electron flow to assess maximal mitochondrial respiration as well as peroxide production. There was no significant effect of treatment on maximum mitochondrial respiration in either permeabilized fibers or heart homogenate (Fig 5 A & B), however treatment with ELAM did decrease amplex red signal in permeablized GAS fibers in forward electron flow but not in heart homogenate (Fig 5 C & D). There was no effect on respiration in either tissue during reverse electron flow (Supplemental Fig 1 A & B), but there was a significant decrease in amplex red signal produced by permeabilized GAS fibers in reverse electron flow using mixed-effect ANOVA (p<0.002) but not in heart (Supplemental Fig 1 C & D).,. It should be noted that total amplex red signal includes not only reactive oxygen species generated by the electron transport chain but also a large portion by lipid peroxides(*17*).

**Figure 5.**
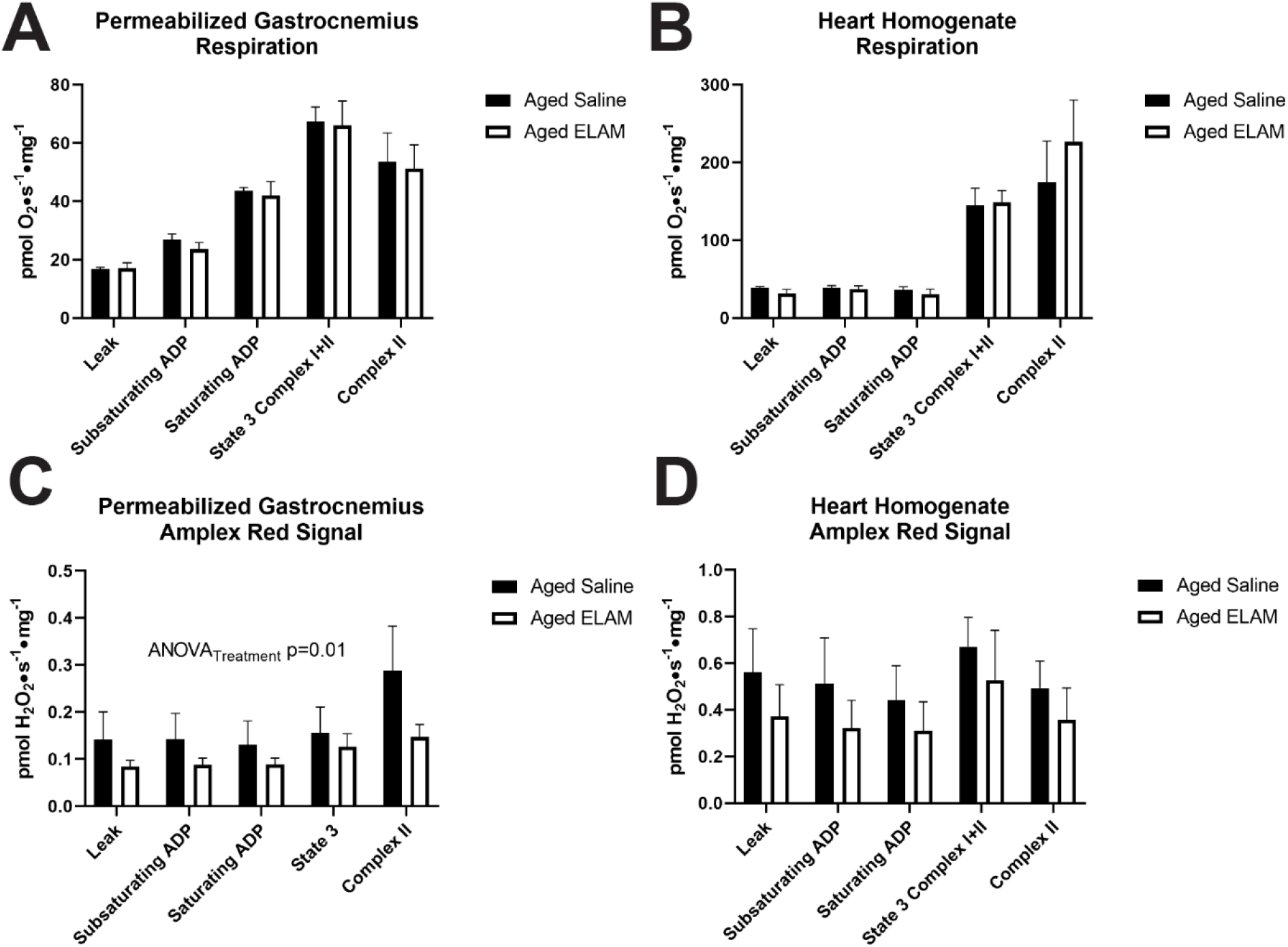
Mitochondrial respiration and amplex red signal. **(A)** Respiration in permeabilized gastrocnemius fibers in forward flow **(B)** Respiration in heart homogenate in forward flow **(C)** Peroxide production in permeabilized gastrocnemius fibers in forward flow **(D)** Peroxide production in heart homogenate in forward flow. Data expressed as means ± SE. Mixed effects ANOVA, with Sidak’s correction for multiple comparisons, p<0.05.

### 3.5 Kidney Pathology

Kidneys are susceptible to predictable age-accumulated injury in both humans and rodents (*18*). With age, kidney function declines and this is accompanied by pathological changes to the microanatomy, most notably focal accumulation of sclerotic lesions in the glomeruli (*19, 20*) and nephron loss (*21*). Loss of specialized glomerular epithelial cells, podocytes, are thought to precede and instigate much of the glomerular dysfunction accumulated with age (*22, 23*). Podocytes are non-proliferative cells that are required for formation of the kidney filtration slit and thus, are paramount to kidney function.

Previously, we demonstrated that eight weeks of continuous systemic ELAM treatment in male mice treated from 24- to 26-months of age was beneficial to the kidney by significantly reducing the accumulation of age-induced glomerulosclerosis (*8*). Additionally, some standard measures of podocyte damage were significantly reduced, suggesting cell-specific protection. In the same study, 28-month-old female mice treated with ELAM for the same 8-week duration exhibited reduced cellular senescence in all kidney compartments. In that study female kidneys were not also assessed for glomerulosclerosis, therefore we sought to determine whether intermittent extended treatment could similarly protect aged female mice from age-accumulated sclerosis. Kidneys were harvested at sacrifice and stained to visualize and quantify microanatomy. Although indicators of kidney aging were observed in all mice, scoring for glomerular injury did not demonstrate any differences between intermittent ELAM injection and untreated female mice (Fig 6 A & B). Further analysis of podocytes for cell number (podocytes per glomerular tuft), cellular hypertrophy (nuclear diameter), cell density (cell number per glomerular tuft area) and glomerular hypertrophy showed no response to ELAM as compared to control animals (Supplemental Figure 2C-F). Thus, kidney improvements, which were detected in male mice with 8-week continuous ELAM intervention were not observed with intermittent intervention in female mice.

**Figure 6.**
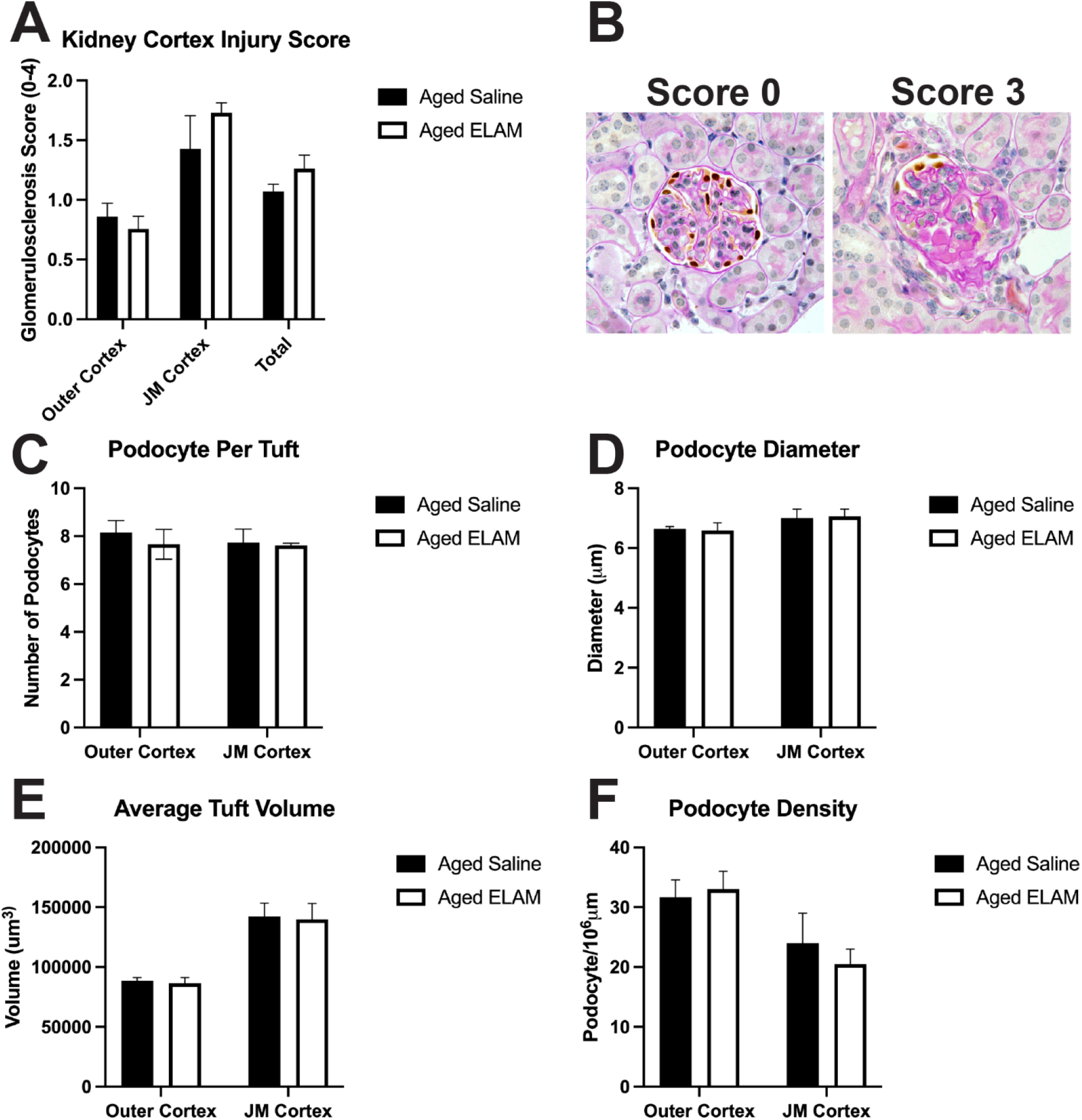
kidney pathology. **(A)** Kidney glomeruli were scored for aging glomerulosclerosis. Because glomerular size differs by kidney region, glomeruli were separated for scoring by cortical location as either occupying the outer cortex or that near the medulla (juxtamedullary/JM). **(B)** Example images of immunohistochemistry of p57 to identify podocyte nuclei (dark brown, DAB) and counter stained with periodic acid Schiff (PAS) and hematoxylin (blue nuclei). Relative scores for example images shown. **(C)** Average number of podocytes per individual glomerular tuft. **(D)** Average diameter of podocyte nuclei within all glomeruli analyzed within each mouse. **(E)** Average volume of glomerular tuft. **(F)** Average density of podocytes within all glomeruli scored for each compartment calculated as number of podocytes normalized to tuft volume. All Control and ELAM treated groups were analyzed by individual Student’s t-test and no significant differences were detected.

## 4. Discussion

This study was designed as a pilot to test whether long-term, intermittent treatment with the mitochondrial targeted peptide ELAM would slow the decline in skeletal muscle and cardiac function when started early in the aging process. We have previously demonstrated that continuous infusion with ELAM using osmotic minipumps for 8 weeks improves skeletal muscle fatigue resistance and *in vivo* mitochondrial function (*6*), reverses cardiac diastolic dysfunction (*7*), and improve tolerance in mouse models of aging when administered late in life (*5*). Furthermore, ELAM has also been demonstrated to be effective in multiple models of chronic disease (*8, 10, 24*) making it a promising mitochondrial targeted intervention for slowing the aging process and reducing frailty. Due to its peptide structure ELAM is rapidly degraded in the gut environment. ELAM has a short elimination half-life of 2-4 hours (*25*), so previous preclinical work with ELAM has focused primarily on daily administration (*5, 9*) or continuous delivery by osmotic minipumps (*6-8*). These approaches are limiting for long-term studies due to the extensive handling of the animals necessary for daily injections, or eye drops administration (*9*), or the multiple surgeries that are required to remove and replace osmotic pumps to maintain delivery for several months. This limitation makes it difficult to compare to other aging interventions such as those in the National Institute of Aging Intervention Testing Program (ITP). Recent evidence in older humans demonstrated that a single treatment with ELAM improved *in vivo* mitochondrial function immediately after treatment and returned to placebo level a week later (*11*). However, this study suggested that effects on skeletal muscle fatigue resistance were elevated a week after treatment. In an attempt to test whether we could reduce the necessary animal handling required for a long-term study we tested whether twice a week treatment with ELAM staring early in life would be sufficient to reproduce or improve our previous results with 8-week treatment on aging cardiac and skeletal muscle function.

Intermittent treatment with ELAM preserved exercise tolerance and delayed decline in skeletal muscle force and cardiac function, but did not lead to improvements in skeletal muscle fatiguability or cardiac function as demonstrated with 8 weeks of continuous delivery (*6, 7*). However, the preserved exercise tolerance observed starting at at 4 weeks (the first time point after baseline) is consistent with the rapid effects of ELAM treatment on mitochondrial function and whole-body performance previously demonstrated with 7 days of daily treatment (*5*). Despite the improved systemic effect on treadmill running, the intermittent treatment did not appear to prevent the loss of body or skeletal muscle mass and morbidity over this age range.

Treadmill exercise tolerance is a complex phenotype involving aspects of motivation as well as physiological capacity. The main physiological capacities that contribute to exercise tolerance are cardiac function and muscle fatigue resistance (*26-28*), both of which are improved with continuous ELAM treatment (*6, 7*). The absence of an effect of ELAM on muscle fatigue resistance of the GAS was surprising given our previous work, although the absence of an improvement in muscle force production is consistent with other studies. Continuous pump treatment for 8 weeks and 4 months of daily injections, both starting at 24 months in C57Bl/6 mice failed to increase muscle force production, despite improvement in mitochondrial function or redox stress. It is now clear that the loss of muscle mass and force production is initiated by the decline in neuromuscular junctions well before significant muscle deficits are observed (*29*). The disruption of NMJ leads to elevated mitochondrial oxidative stress in the myofiber that leads to a feedforward cycle causing further damage to the NMJ (*30, 31*). Although the ability of ELAM to reduce mitochondrial oxidative stress is well-established, the twice per week treatment used in the current study, even when initiated early in the process, does not appear to be sufficient to prevent this cycle and preserve skeletal muscle force production.

The primary aging cardiac phenotypes in the C57Bl/6 mice are diastolic dysfunction and cardiac hypertrophy. Previous work demonstrated that ELAM reversed diastolic dysfunction as assessed by the E’/A’ ratio with 8 weeks of continuous treatment (*7*) and this effect persisted for two weeks following the end of treatment(*7*). In contrast, the intermittent treatment did not demonstrate any effect on diastolic dysfunction until 28 weeks of treatment. There are two potential explanations for the greater length of treatment required in this study. The first is that the intermittent treatment required a longer period to drive the cardiac remodeling underlying improved cardiac relaxation and the persistent effect of treatment (*7*). The alternative is that there was not significant cardiac dysfunction present until later in the treatment period. Multiple studies have demonstrated that ELAM has no effect on healthy, well-functioning heart. This interpretation is supported by the stable E’/A’ ratio over the first 24 weeks of this study with values similar to what has been reported for young C57Bl/6 mice. The ability of intermittent treatment to contribute to partial remodeling of the aged heart is supported by the effect on LVMI, although this did not translate to reduced cardiac hypertrophy based on total heart mass.

## 5. Conclusions

This pilot study demonstrated that, although twice weekly treatment with ELAM for 8 months led to preservation of whole-body exercise tolerance and partial improvement of cardiac aging phenotypes, it only reproduced a portion of the significant benefits previously observed on skeletal and cardiac function with continuous 8-week treatment at late age. Thus, continuous ELAM treatment (daily injection or osmotic pump delivery) remains the most effective delivery method to observe long-term benefits of ELAM on lifespan and healthspan in aging mice.

## Abbreviations

GAS: Gastrocnemius
TA: Tibialis Anterior
ELAM: elamipretide

## Statements and Declarations

The authors declare no competing financial interests.

## 6. Acknowledgements

The authors would like to thank Rudy Stuppard for technical assistance with all aspects of this study. elamipretide was provided by Stealth Biotherapeutics, Inc.

## 10. Supplement

**Supplemental Figure 1.**
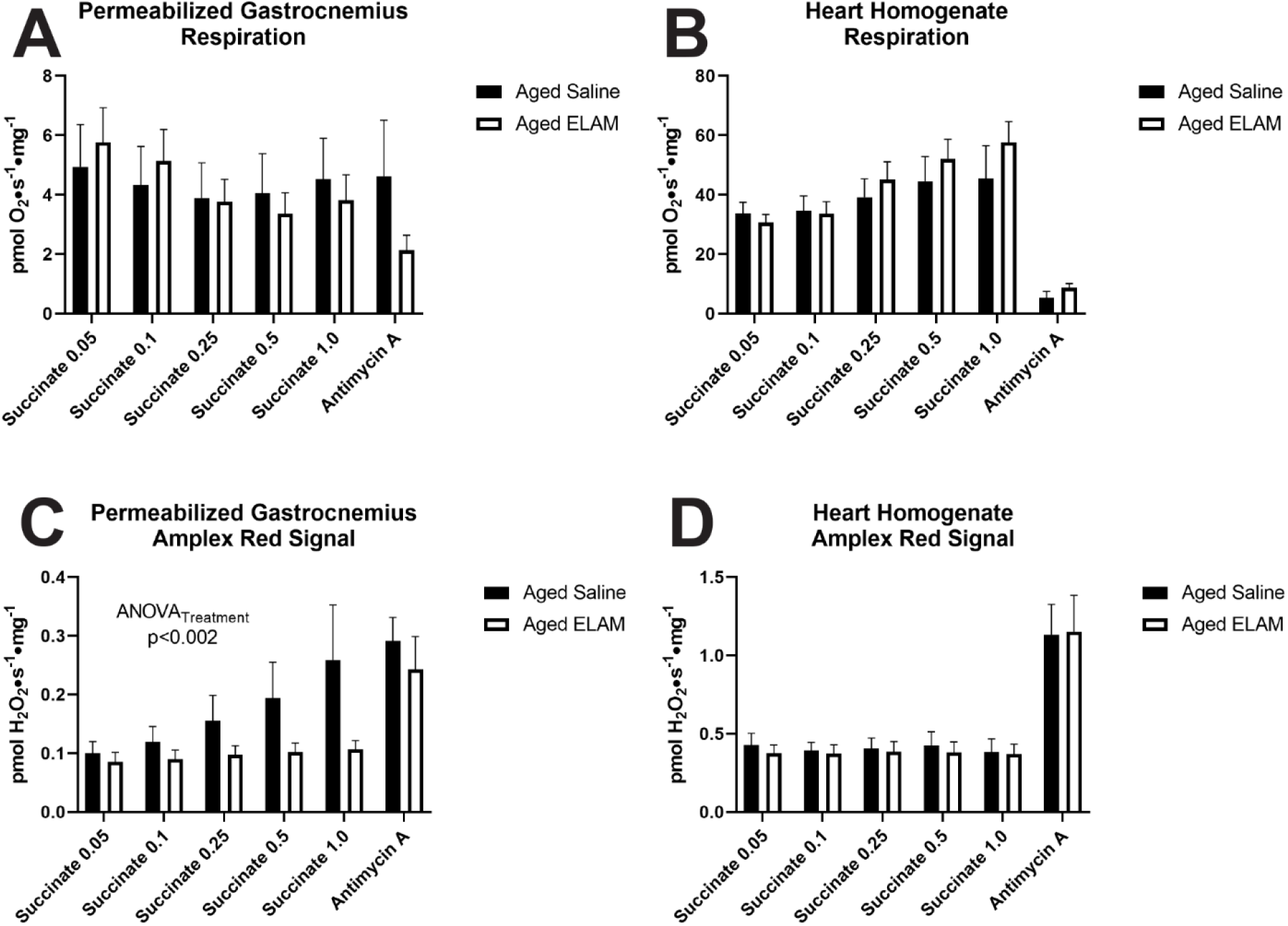
Mitochondrial respiration and amplex red signal. **(A)** Peroxide production in permeabilized gastrocnemius fibers in reverse flow **(B)** Peroxide production in heart homogenate in reverse flow **(C)** Respiration in permeabilized gastrocnemius fibers in reverse flow **(D)** Respiration in heart homogenate in reverse flow. Data expressed as means ± SE. Mixed effects ANOVA, with Sidak’s correction for multiple comparisons, p<0.05.

## References

1. Q. Tian et al., Muscle mitochondrial energetics predicts mobility decline in well-functioning older adults: The baltimore longitudinal study of aging. Aging Cell 21, e13552 (2022).

2. S. C. Lewsey et al., Exercise intolerance and rapid skeletal muscle energetic decline in human age-associated frailty. JCI Insight 5, (2020).

3. N. Sun, R. J. Youle, T. Finkel, The Mitochondrial Basis of Aging. Mol Cell 61, 654–666 (2016).

4. T. E. S. Kauppila, J. H. K. Kauppila, N. G. Larsson, Mammalian Mitochondria and Aging: An Update. Cell Metab 25, 57–71 (2017).

5. M. P. Siegel et al., Mitochondrial-targeted peptide rapidly improves mitochondrial energetics and skeletal muscle performance in aged mice. Aging Cell 12, 763–771 (2013).

6. M. D. Campbell et al., Improving mitochondrial function with SS-31 reverses age-related redox stress and improves exercise tolerance in aged mice. Free Radic Biol Med 134, 268–281 (2019).

7. Y. A. Chiao et al., Late-life restoration of mitochondrial function reverses cardiac dysfunction in old mice. Elife 9, (2020).

8. M. T. Sweetwyne et al., The mitochondrial-targeted peptide, SS-31, improves glomerular architecture in mice of advanced age. Kidney Int 91, 1126–1145 (2017).

9. N. M. Alam, R. M. Douglas, G. T. Prusky, Treatment of age-related visual impairment with a peptide acting on mitochondria. Dis Model Mech 15, (2022).

10. S. Tarantini et al., Treatment with the mitochondrial-targeted antioxidant peptide SS-31 rescues neurovascular coupling responses and cerebrovascular endothelial function and improves cognition in aged mice. Aging Cell 17, (2018).

11. B. Roshanravan et al., In vivo mitochondrial ATP production is improved in older adult skeletal muscle after a single dose of elamipretide in a randomized trial. PLoS One 16, e0253849 (2021).

12. P. Jousilahti, E. Vartiainen, J. Tuomilehto, P. Puska, Sex, age, cardiovascular risk factors, and coronary heart disease: a prospective follow-up study of 14 786 middle-aged men and women in Finland. Circulation 99, 1165–1172 (1999).

13. B. J. North, D. A. Sinclair, The intersection between aging and cardiovascular disease. Circ Res 110, 1097–1108 (2012).

14. M. C. White et al., Age and cancer risk: a potentially modifiable relationship. Am J Prev Med 46, S7–15 (2014).

15. V. Therakomen, A. Petchlorlian, N. Lakananurak, Prevalence and risk factors of primary sarcopenia in community-dwelling outpatient elderly: a cross-sectional study. Sci Rep 10, 19551 (2020).

16. M. Venkatareddy et al., Estimating podocyte number and density using a single histologic section. J Am Soc Nephrol 25, 1118–1129 (2014).

17. G. Pharaoh et al., Targeting cPLA2 derived lipid hydroperoxides as a potential intervention for sarcopenia. Sci Rep 10, 13968 (2020).

18. E. D. O’Sullivan, J. Hughes, D. A. Ferenbach, Renal Aging: Causes and Consequences. J Am Soc Nephrol 28, 407–420 (2017).

19. A. Denic, R. J. Glassock, A. D. Rule, Structural and Functional Changes With the Aging Kidney. Adv Chronic Kidney Dis 23, 19–28 (2016).

20. J. B. Hodgin et al., Glomerular Aging and Focal Global Glomerulosclerosis: A Podometric Perspective. J Am Soc Nephrol 26, 3162–3178 (2015).

21. A. Denic et al., The Substantial Loss of Nephrons in Healthy Human Kidneys with Aging. J Am Soc Nephrol 28, 313–320 (2017).

22. J. Wiggins, Podocytes and glomerular function with aging. Semin Nephrol 29, 587–593 (2009).

23. M. Camici, A. Carpi, G. Cini, F. Galetta, N. Abraham, Podocyte dysfunction in aging--related glomerulosclerosis. Front Biosci (Schol Ed) 3, 995–1006 (2011).

24. W. Zhao et al., Elamipretide (SS-31) improves mitochondrial dysfunction, synaptic and memory impairment induced by lipopolysaccharide in mice. J Neuroinflammation 16, 230 (2019).

25. H. H. Szeto, A. V. Birk, Serendipity and the discovery of novel compounds that restore mitochondrial plasticity. Clin Pharmacol Ther 96, 672–683 (2014).

26. J. J. Wan, Z. Qin, P. Y. Wang, Y. Sun, X. Liu, Muscle fatigue: general understanding and treatment. Exp Mol Med 49, e384 (2017).

27. M. A. Nystoriak, A. Bhatnagar, Cardiovascular Effects and Benefits of Exercise. Front Cardiovasc Med 5, 135 (2018).

28. K. Albouaini, M. Egred, A. Alahmar, D. J. Wright, Cardiopulmonary exercise testing and its application. Postgrad Med J 83, 675–682 (2007).

29. G. K. Sakellariou et al., Neuron-specific expression of CuZnSOD prevents the loss of muscle mass and function that occurs in homozygous CuZnSOD-knockout mice. FASEB J 28, 1666–1681 (2014).

30. Y. C. Jang et al., Increased superoxide in vivo accelerates age-associated muscle atrophy through mitochondrial dysfunction and neuromuscular junction degeneration. FASEB J 24, 1376–1390 (2010).

31. F. L. Muller et al., Absence of CuZn superoxide dismutase leads to elevated oxidative stress and acceleration of age-dependent skeletal muscle atrophy. Free Radic Biol Med 40, 1993–2004 (2006).

